# Optimizing population simulations to accurately parallel empirical data for digital breeding

**DOI:** 10.1101/2025.06.18.660215

**Authors:** Michael J. Burns, Rafael Della Coletta, Samuel B. Fernandes, Martin O. Bohn, Alexander E. Lipka, Candice N. Hirsch

**Author notes:** authors contributed equally to this work (Michael J. Burns, Rafael Della Coletta).

## Abstract

The use of computational and data-driven approaches to accelerate and optimize breeding programs is becoming common practice among plant breeders. Simulations allow breeders to evaluate potential changes in breeding schemes in a time– and cost-efficient manner. However, accurately simulating traits that match empirical trait data remains a challenge. Here we tested if incorporating information about the genetic architecture from genome-wide association studies (GWAS) of maize agronomic traits with varying heritabilities into simulations can improve the concordance between simulated and empirical data in a population of hybrids developed from crosses of 333 maize recombinant inbred lines grown in four to eleven environments. Using at least 200 non-redundant top GWAS hits as causative variants, regardless of statistical significance, resulted in mean correlations between simulated and empirical trait data of 0.397 to 0.616 within environments and 0.610 to 0.915 across environments. Reducing the GWAS estimated marker effect sizes in the simulations further improved concordance with empirical data. This study provides valuable insights into methods for simulating more realistic phenotypes for digital breeding to parallel empirical trait distributions, and that these simulated traits are highly concordant with observed variance partitioning (i.e. genotype, environment, etc.), and genomic prediction performance.

**PLAIN LANGUAGE SUMMARY:** Plant breeders need to evaluate many possible changes to breeding schemes and resource allocation. One method of optimizing these changes is through computer-based simulations, which can be time– and cost-efficient. The genetic architecture of randomly simulated traits rarely parallels the genetic architecture of real-world traits, which can mislead the interpretation of such simulations. In this study, the genetic architecture of real-world traits was used to inform the simulation of digital traits to develop an optimized simulation pipeline for creating and assessing simulated breeding programs. The utility of these informed simulations is demonstrated through genomic prediction assessment, in which we show that the rank order of individuals and the distribution of traits is similar between simulated and real-world data.

**CORE IDEAS:**

● Simulations allow breeders to evaluate potential changes in breeding schemes in a time– and cost-efficient manner.
● Incorporating the genetic architecture of traits in simulations can improve the concordance with empirical data.
● Reducing GWAS marker effect size improves correlation and distribution concordance of simulated and empirical data.
● Increasing the number of causative variants beyond GWAS significance thresholds improves simulation performance.
● Simulated data used in genomic prediction produces results similar to empirical genomic predicted data.

## 1 INTRODUCTION

Resource allocation is one of the biggest decision points in a breeding program. Simulations are a valuable tool for advancing plant breeding by providing a fast and inexpensive way to test hypotheses, evaluate new breeding schemes, and identify potential problems before moving forward with costly and time-consuming experiments (X. Li et al., 2012). For example, simulation studies formulated the foundation and usefulness of genomic prediction and selection in plant and animal breeding by showing the influence of population size and relatedness, heritability, and number of markers on prediction accuracy (Combs & Bernardo, 2013; Daetwyler et al., 2010, 2013; Meuwissen et al., 2001).

Simulating phenotypes with genetic architectures that match reality is key for translating knowledge from simulations to real life when running digital and empirical breeding programs in parallel. Currently, there are many available tools for simulating phenotypes, with varying applicability to different scenarios (Gaynor et al., 2021; Shrote & Thompson, 2024; Sun et al., 2011; Werner et al., 2024; Younis et al., 2023; L. Zhang et al., 2022). For example, AlphaSim is a software tool that enables comprehensive simulation of an entire breeding program (Faux et al., 2016; Gaynor et al., 2021). It allows backwards– and forwards-in-time simulations of haplotypes and phenotypes, testing of different crossing and selection schemes, and modeling of genomic prediction throughout the simulated breeding program (Faux et al., 2016; Gaynor et al., 2021). PhenotypeSimulator is another powerful software that focuses on simulating multiple traits with complex multi-locus genetics, including genetic variant and background (infinitesimal) effects, non-genetic effects, and noise effects with different covariance structures (Meyer & Birney, 2018). However, despite offering a lot of flexibility on multiple stages of simulations, these programs do not allow easy incorporation of real-world data, such as effect sizes based on genotype-phenotype association studies or environmental correlations from target breeding environments. The R package simplePHENOTYPES was developed to provide users with the ability to control every aspect of trait genetic architecture when simulating complex multi-trait phenotypes, including pleiotropy, based on real genotypic and environmental information (Fernandes & Lipka, 2020). This flexibility is important as it allows users to optimize parameter combinations that result in simulations that approach reality. It also allows simulations of traits across multiple environments by providing an environmental correlation matrix while keeping the same causative variants across these environments with varying effect sizes in each environment.

To input parameters that mirror reality, it is necessary to understand the genetic architectures underlying empirical traits. Genome-wide association studies (GWAS) and quantitative trait loci (QTL) studies are both methods used to identify genomic regions that contribute to phenotypic variation and the effect size of the region (Tibbs Cortes et al., 2021). QTL studies are commonly used in experimental populations (such as biparental crosses), whereas GWAS are performed on a larger and more diverse population, allowing for the detection of regions with smaller effect sizes at higher resolution (Tibbs Cortes et al., 2021). GWAS has been used extensively in plant breeding to identify the factors underlying the genetic architecture of many different traits in a wide range of crops (H.-J. Liu & Yan, 2019). In some cases, the genetic markers with the most significant p-values for phenotypes have been useful to increase the accuracy of genomic prediction when incorporated into the models (Spindel et al., 2016; Z. Zhang et al., 2014). However, this is not always the case (Rice & Lipka, 2019), and the importance of incorporating such information into simulations of phenotypes has yet to be documented.

In this study, the utility of different methods for incorporating the genetic architecture of empirical traits into trait simulation was tested by accounting for modes of gene action, modifying the number of included causative markers, and adjusting the effect size of the included markers. To test how well the simulated data mirrored the empirical data, performance metrics including the Spearman Rank correlation coefficient, Kolmogorov-Smirnov D-statistic, t-test statistic, and log of the F-test f-ratio were used to monitor shifts in trait means and distributions when simulation parameters changed. Finally, genomic prediction was performed on the empirical and simulated data. The genomic prediction of simulated data was compared to the simulated data to assess the utility as a digital twin, and to the genomic prediction of empirical data to assess drift across subsequent levels of inference.

## 2 MATERIALS AND METHODS

### 2.1 Phenotypic data

A set of 333 F_7_ recombinant inbred lines (RILs) from half diallel crosses among six ex-PVP maize inbred lines (B73, PHG39, PHG47, PH207, PHG35, and LH82; Della Coletta et al., 2023) were crossed to produce 400 F_1_ hybrids (Table S1). These hybrids were grown in six locations across the United States Midwest in 2019 and/or 2020 for a total of 11 environments (i.e., Location × Year combination; Table S2). In each environment two replications were planted in a randomized complete block design. Phenotypic measurements were obtained for ear height, plant height, grain moisture, and grain yield in 4-11 of the environments for 352-400 of the hybrids per environment (Table S3). Ear height (cm) and plant height (cm) were manually measured as the height from the ground to the ear bearing node and from the ground to the ligule of the flag leaf. Grain moisture (%) and grain yield (bu ac^-1^) were obtained from plot combine measurements and grain yield was normalized to 15.5% moisture. For each trait, a linear mixed model was fit with the R package lme4 (Bates et al., 2015) to evaluate the percent of variance explained by genotype, environment, and genotype-by-environment effects using the linear model:

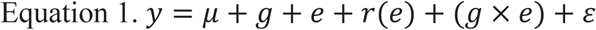

where *y* is the trait values, μ is the overall mean, *g* is the fixed genotypic effect of a maize hybrid, *e* is the random environmental effect, *r*(*e*) is the random replicate effect nested within environment, *g* × *e* is the random effect of genotype-by-environment interactions, and ε is the residual effect with variance 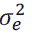 with 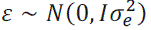. Narrow sense heritability for each trait in each environment was determined using equation 2 to model genetic and non-genetic effects, where *y* is the trait values, μ is the overall mean, *g* is the random genotypic effect, *r* is the random replicate effect, and ε is the residual effect with variance 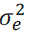 with 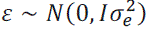.

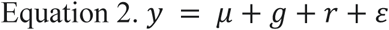

The variance components of equation 2 were used to calculate narrow sense heritability with the following equation:

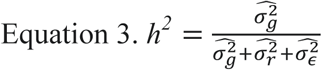

where 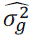 is the estimated genetic variance, 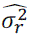 is the estimated replication variance, and 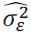 is the residual variance.

### 2.2 Genotypic data

Genomic variants were previously identified for the six inbred parents of this population, including single nucleotide polymorphisms (SNPs; Mazaheri et al., 2019) and structural variants (SVs; Della Coletta et al., 2023), and these variants were previously projected to the 333 RILs (Della Coletta et al., 2023). Using this dataset, genotypic data were generated *in silico* for each hybrid that was empirically tested in this study by combining the alleles of its respective RIL parents at each locus. The hybrid genotype was set to missing data if at least one of its parents had a heterozygous genotype, as it cannot be known which allele was passed on to the hybrid. The hybrid genotypic dataset contained 8,510,449 SNPs and 8,416 SVs with a mean of 41.4% missing data per genotype and a mean of 13.5% missing data per marker.

### 2.3 Environmental data

Prior to simulating traits across multiple environments, a weather-based correlation matrix among the 11 growth environments of this study was obtained using the R package EnvRtype v1.1.0 (Costa-Neto et al., 2021). For this matrix, 17 weather parameters (Table S4) for each environment in 10-day intervals throughout the season (April 1st to October 31st) were used (Table S5). The weather data was transformed into a variance-covariance environmental relatedness matrix with the env_kernel function in EnvRtype v1.1.0 (Costa-Neto et al., 2021). The final correlation matrix among environments (Table S6) was generated using the cov2cor function in R v3.6.0 (R Core Team, 2019).

### 2.4 Genetic architecture of phenotypic traits

To determine the genetic architecture underlying each of the four empirically measured traits (i.e., ear height, plant height, grain moisture, and grain yield), genome-wide association studies (GWAS) were performed using a unified mixed linear model (MLM; Yu et al., 2006) with additive or dominance effects implemented in GAPIT v3 (Wang & Zhang, 2021).

To this end, best linear unbiased estimates (BLUEs) were calculated for each hybrid in each environment with Equation 4.

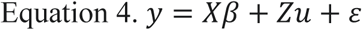

where *y* is the vector of phenotypic values in one environment, *X* is the design matrix for the fixed genotype effects β, *Z* is the design matrix for the random replication effects *u* with 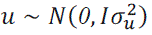, *I* is an identity matrix, 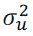 is the replication variance, and ε is the residual effect with variance 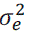 with 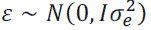. The BLUEs were calculated in ASReml-R v4.1 (Butler et al., 2017).

Prior to conducting the GWAS, the genotypic dataset was filtered to remove markers in high linkage disequilibrium (LD; r^2^ > 0.9) within 10 kb windows, with minor allele frequency lower than 5%, and/or missing data rate higher than 25% using PLINK v1.9 (Chang et al., 2015). The window size for LD pruning was defined based on the rapid LD decay of the inbred parents (Della Coletta et al., 2023) and to maximize the number of both SNP and SV markers after filtering. The filtered hybrid genotypic dataset contained 359,990 SNPs and 3,110 SVs. The filtered matrix was then converted into two different numeric format matrices for additive and dominance models using GAPIT v3 (Wang & Zhang, 2021). For additive GWAS models, homozygous genotypes were encoded as either 0 or 2 and heterozygous calls were encoded as 1. For dominance GWAS models, all homozygous calls were encoded as 0 and heterozygous calls were encoded as 1. Principal component analysis (PCA) was performed with the GAPIT.PCA function in GAPIT v3.4 (Wang & Zhang, 2021) for input into the GAPIT function with model set to MLM. It was determined that most of the genetic variation in the population could be explained by the first seven principal components (PCs). These seven PCs were used as covariates in the GWAS models to control for population structure. A genetic kinship matrix was obtained using the VanRaden method (VanRaden, 2008) implemented in GAPIT v3.4 (Wang & Zhang, 2021).

The statistical significance of a marker associated with a phenotype was determined by permutations based on the FarmCPU.P.Threshold function from FarmCPU v1.02 (X. Liu et al., 2016). This function was adapted to accommodate the MLM model within GAPIT. Briefly, BLUEs were shuffled 30 times, and the lowest p-value between a marker and a phenotype was recorded each time. The final p-value threshold to be used in a GWAS for each trait was then determined based on the 95% quantile of permuted p-values. Finally, the phenotypic variance explained (PVE) was calculated for each trait using all available markers and using only the significant GWAS markers plus the 100 non-redundant markers with the lowest p-value using the software GCTA v1.94 (Yang et al., 2011). The phenotypic input files for this analysis were the BLUE values of each trait in each environment for each hybrid.

### 2.5 Variant selection for simulations

Multiple genetic markers can have a high association with the same causative variant, and if linked, can account for the same variation. To determine a set of non-redundant markers for simulations, LD was calculated among all markers using PLINK v1.9 (Chang et al., 2015). The window size for this calculation was set to the chromosome size to obtain LD values between markers in the same chromosome, but not across chromosomes. For each trait and GWAS model, the markers in each environment were ranked based on their p-value in that environment, and the mean rank across environments was calculated. The top 5, 10, 20, 30, 40, 50, 60, 70, 80, 90, 100, 150, 200, 250, 300, 350 mean ranked markers were then selected as causative variants to observe where concordance between simulated and empirical data plateaus. Each time a marker was selected, any markers in high LD (LD>0.9) with that marker were removed from the set of remaining markers to reduce redundancy.

### 2.6 Trait simulation across multiple environments

Multi-environment phenotypic values were simulated for the hybrid population described above based on the genetic architecture identified by GWAS for each trait using the R package simplePHENOTYPES v1.3 (Fernandes & Lipka, 2020). This package allows the simulation of traits based on real genotypic data, where the user can select which markers are the causative variants and define their effect sizes and types (i.e., additive or dominance). Traits can be simulated across multiple environments by repeating the simulations with the same causative variants but changing their effect sizes and adding a correlation matrix that defines the relationship among the environments.

For each simulated trait, the heritability and means from the corresponding empirical trait data were used as inputs into simplePHENOTYPES. The causative variants were selected based on different numbers of top non-redundant additive and dominance GWAS markers across environments (e.g., 5, 10, 20, etc.). The gene mode of action of these markers was set to additive or dominant according to the GWAS model that was used to identify the marker as a causative variant. The estimated effect size of each marker was first set to the value from the GWAS within each environment. The simulations were repeated a second time with a reduced effect size, where the effect of each marker was reduced to 10% of its original GWAS effect size.

The total number of environments simulated also matches that of the empirical data (i.e., 4 environments for ear height, 5 for plant height, 11 for grain moisture, and 10 for grain yield), and the empirical environmental correlation matrix was used as the residual correlation matrix in simplePHENOTYPES. Two replicates per hybrid were simulated in each environment. If replicates were not correlated within environments (Pearson’s correlation < 0.1 in any environment), a third replicate was created and used to replace the second replicate.

An ANOVA was performed for each simulated trait using equation 1 to determine the percent variance explained by each experimental factor. Spearman Rank correlation coefficient (cor; Spearman, 1904), Kolmogorov-Smirnov D-statistic (ks.test; Simard & L’Ecuyer, 2011), t-test statistic (t.test; Student, 1908), and log of the F-test statistic (var.test; Box, 1953) were collected in R v4.3.0 (R Core Team, 2019) to measure the concordance between simulated and empirical phenotypic datasets. These metrics demonstrated whether similar trends across and within environments were observed and how the mean and shape of trait distributions changed after modifying simulation parameters.

### 2.7 Genomic prediction models

Genomic prediction models were run for simulated and empirical trait data in two stages. First, best linear unbiased estimates (BLUEs) were calculated for each environment to account for genotype-by-environment effects using the following mixed linear model implemented in ASReml-R v4.1 (Butler et al., 2017):

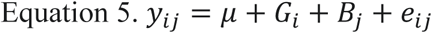

where *y*_*+_ is the trait value of hybrid *i* in replicate *j*, μ is a constant, *G*_*_ is the fixed effect of the *i*th hybrid, *B*_+_ is the random effect of the *j*th replicate with 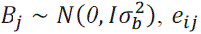 is random effect of residuals with 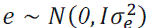, and *I* is an identity matrix.

In the second stage, a random set of 500 markers from the pruned genotypic dataset were selected as the predictors for the genomic prediction models. This process of selecting 500 random markers was repeated ten times to minimize random sampling bias. Multivariate genomic best linear unbiased prediction (GBLUP) models were run with additive or dominance relationships with each set of 500 random predictors using the following mixed linear model also implemented in ASReml-R v4.1 (Butler et al., 2017):

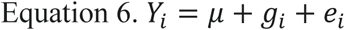

where *Y*_*_ is the vector of multivariate responses associated with hybrid *i* (*i* = *1,2*, …, *n*) in which *Y*_*i*_ = [*Y*_*i1*_, *Y*_*i2*_, …, *Y*_*ip*_]^T.^ and *p* is the number of environments, μ is the vector of the constants associated with each environment, *g*_*_ is the vector of random effects of hybrid *i* associated with each environment with *g* ∼ *N_np_*(*0*, *G* ⊗ *A*) or *g* ∼ *N_np_*(*0*, *G* ⊗ *D*) where *G* is the variance-covariance matrix for genetic effects, *A* is the realized additive relationship matrix and *D* is the realized dominance relationship matrix, *e*_*_ is the vector of random effects of residuals with *e* ∼ *N_np_*(*0*, *I* ⊗ *R*) with *R* as the variance-covariance matrix for residual effects, and *I* as an identity matrix. The VanRaden method (VanRaden, 2008) was used to calculate the additive relationship matrix, while the dominance relationship matrix was calculated using the Vitezica method (Vitezica et al., 2013). Both matrices were calculated from genotypic data using the Gmatrix function of the R package AGHmatrix (Amadeu et al., 2016). The *G* matrix was modeled with a first-order factor analytic structure where the genetic variance and covariance were allowed to be different among environments, while the *R* matrix was diagonal (i.e., residual variance was different among environments, and covariance among environments was not allowed to be different). For computational efficiency, the total number of environments with grain moisture and grain yield data was reduced to six by removing highly correlated environments (BAY19, BEC-BL19, BEC-BL20, BEC-EP20, and URB19 for grain moisture, and BEC-BL19, BEC-BL20, BEC-EP20, and URB19 for grain yield).

Prediction accuracy was computed using the two cross-validation schemes (CV2 and CV1) described in (Burgueño et al., 2012). Briefly, hybrids were divided into five distinct groups, and the phenotypes of hybrids belonging to one group were predicted using information from hybrids in the four other groups. This process was repeated until the hybrids from all groups had their phenotypes predicted. The prediction accuracy was then determined by calculating Pearson’s correlation coefficient between observed and predicted values. The difference between the cross-validation schemes CV2 and CV1 was that, in CV2, hybrids to be predicted still had phenotypic information available in some environments, and in CV1, they did not have any information retained across environments. The cross-validation process was repeated three times to minimize sampling bias when assigning hybrids to their respective groups. Spearman rank correlations between predicted values from simulated trait data and the empirical trait values predicted with the same set of 500 markers were calculated using the cor function in R v4.3.0 (R Core Team, 2019).

## 3 RESULTS AND DISCUSSION

### 3.1 GWAS revealed distinct genetic factors contributing to trait variability across environments

The overall goal of this study was to evaluate the utility of incorporating the genetic architecture of a trait into trait simulations to allow for the development of digital breeding programs that parallel empirical data as closely as possible. To this end, four traits with varying genetic complexity and heritability were evaluated, including: ear height (4 environments), plant height (5 environments), grain moisture (11 environments), and grain yield (10 environments). Each hybrid had BLUEs calculated for each trait in each environment (Figure 1A). The heritabilities for these traits in each environment ranged from 0.21 to 0.94 (Table 1). The variance partitioning showed a high influence of environmental factors affecting grain moisture and yield compared to ear height and plant height (Figure 1B and Supplemental Table S7), similar to other studies in maize (Ndhlela et al., 2014; Peiffer et al., 2014; Renk et al., 2021).

**Figure 1.**
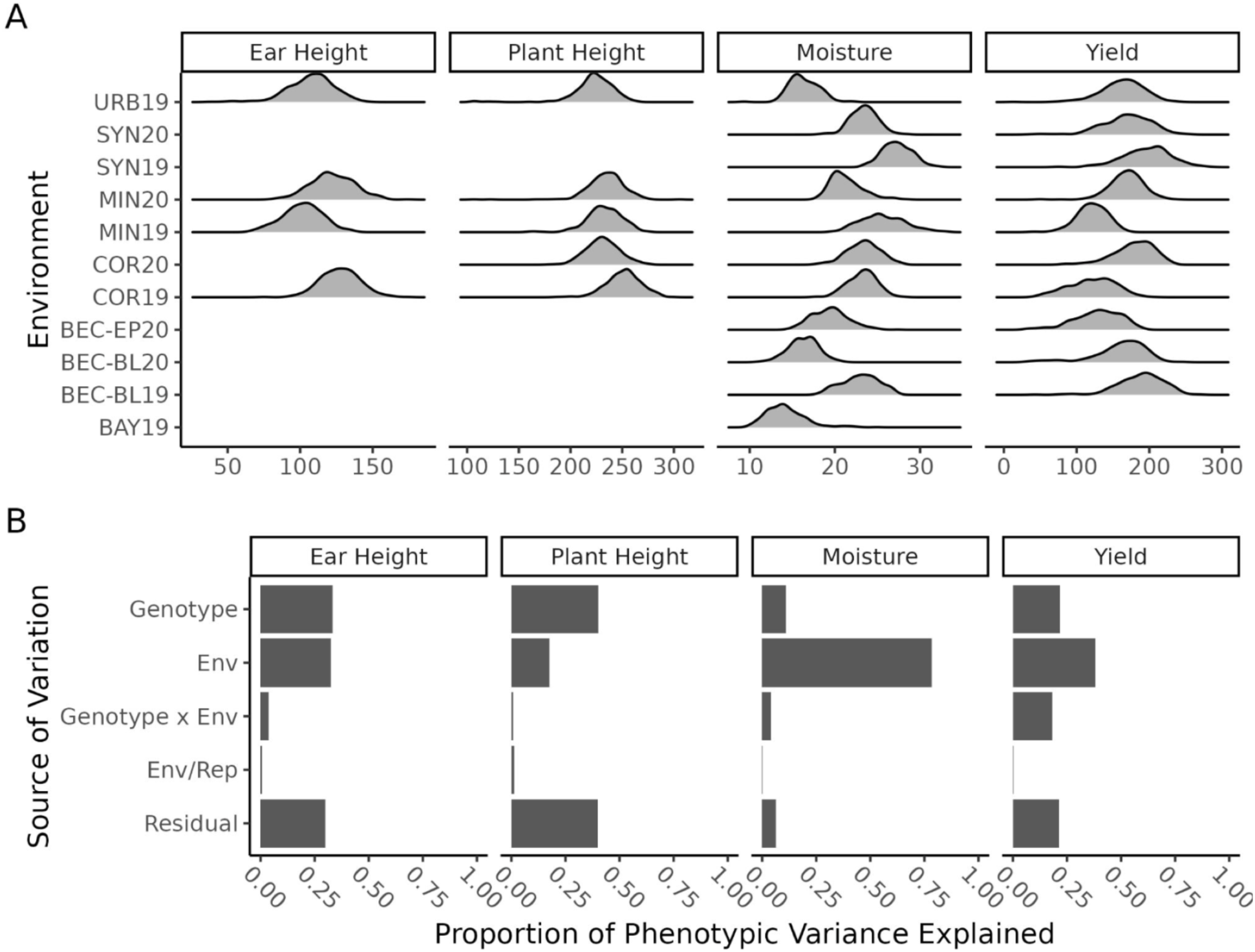
Distribution of traits and their variance partitioning. (A) Distribution of best linear unbiased estimates for ear height (cm), plant height (cm), moisture (%), and yield (bu/ac) in each environment the given trait was recorded in. (B) Proportion of phenotypic variance explained by genotype, environment, genotype-by-environment, replication-within-environment, and residual effects from analysis of variance (ANOVA) for ear height, plant height, grain moisture, and grain yield collected from 354 to 399 hybrids in 4 to 11 environments.

**Table 1.** Summary of phenotypic values and heritabilities for traits in 4 to 11 environments.

| **Trait** | **Environment** | **Mean (± SE)<sup>a</sup>** | **Range<sup>a</sup>** | ***h*<sup>2</sup>** |
| --- | --- | --- | --- | --- |
| Ear Height | COR19 | 127.6 ± 0.7 | 75 - 174 | 0.52 |
| Ear Height | MIN19 | 101.8 ± 0.7 | 67 - 136 | 0.54 |
| Ear Height | MIN20 | 122.2 ± 0.8 | 80 - 174 | 0.63 |
| Ear Height | URB19 | 107.4 ± 0.8 | 38 - 147 | 0.4 |
| Moisture | BAY19 | 14.2 ± 0.1 | 10 - 25 | 0.73 |
| Moisture | BEC-BL19 | 23 ± 0.1 | 18 - 28 | 0.91 |
| Moisture | BEC-BL20 | 16.3 ± 0.1 | 11-21 | 0.88 |
| Moisture | BEC-EP20 | 19.4 ± 0.1 | 14 - 27 | 0.94 |
| Moisture | COR19 | 23.2 ± 0.1 | 17 - 28 | 0.59 |
| Moisture | COR20 | 23.4 ± 0.1 | 18 - 28 | 0.71 |
| Moisture | MIN19 | 25.6 ± 0.1 | 20 - 33 | 0.31 |
| Moisture | MIN20 | 21.3 ± 0.1 | 17 - 29 | 0.76 |
| Moisture | SYN19 | 27.3 ± 0.1 | 23 - 32 | 0.71 |
| Moisture | SYN20 | 23.4 ± 0.1 | 18 - 29 | 0.74 |
| Moisture | URB19 | 16.4 ± 0.1 | 10 - 22 | 0.53 |
| Plant Height | COR19 | 251.3 ± 0.8 | 186 - 298 | 0.71 |
| Plant Height | COR20 | 231.2 ± 0.8 | 179 - 277 | 0.65 |
| Plant Height | MIN19 | 232.7 ± 0.9 | 159 - 275 | 0.52 |
| Plant Height | MIN20 | 234.4 ± 0.9 | 106 - 306 | 0.48 |
| Plant Height | URB19 | 222.8 ± 1.1 | 106 - 265 | 0.21 |
| Yield | BEC-BL19 | 188.7 ± 1.6 | 43 - 271 | 0.67 |
| Yield | BEC-BL20 | 159.7 ± 1.9 | 30 - 247 | 0.8 |
| Yield | BEC-EP20 | 128.8 ± 1.6 | 31 - 190 | 0.88 |
| Yield | COR19 | 121.4 ± 1.7 | 43 - 206 | 0.56 |
| Yield | COR20 | 179 ± 1.5 | 81 - 229 | 0.59 |
| Yield | MIN19 | 121.3 ± 1.1 | 39 - 166 | 0.52 |
| Yield | MIN20 | 168.6 ± 1.2 | 48 - 238 | 0.57 |
| Yield | SYN19 | 196.4 ± 1.7 | 72 - 286 | 0.59 |
| Yield | SYN20 | 169.2 ± 1.7 | 34 - 256 | 0.57 |
| Yield | URB19 | 163.3 ± 1.5 | 13 - 243 | 0.42 |

<sup>a</sup> Ear height and plant height are measured in centimeters, grain moisture is measured as a percentage, and yield is measured in bushels per acre.

Given that there was significant genotypic variation for each of the traits (Supplemental Table S7), the genetic architectures of each trait in each environment were determined. GWAS was performed on the BLUEs of each hybrid calculated for each trait (N=4) in each environment (N=4 to 11) using the unified mixed linear model (MLM) with either additive or dominance marker effects. A total of two significant loci for ear height, four for plant height, 17 for grain moisture, and 22 for grain yield were identified (Supplemental Figure S1, Table S8). Similar numbers of markers associated with these traits were found in other maize studies (W. Li et al., 2021; Mazaheri et al., 2019; Ndlovu et al., 2022; Xue et al., 2013), with five of them being within ∼2 to 3 Mb of the markers in this study (Mazaheri et al., 2019; Ndlovu et al., 2022). Since the MLM can report more than one marker tagging the same causative variant, significant and non-significant markers that were in high LD with each other were removed from downstream analysis, and only the most significant marker was retained.

To better understand how well the significant markers represent the genetic architecture of each trait, the percentage of phenotypic variance explained by the significant markers was evaluated. On average, ∼37% (ranging from 7% to 55%) of phenotypic variance was accounted for when using all ∼360,000 genome-wide markers, while only 1.47% to 10.47% of the variance was explained by the significant markers. Given the low number of significant markers, the percentage variance explained was also evaluated with the top 100 non-significant markers included. The p-value threshold for these 100 non-significant markers ranged from 7.87 x 10^-6^ to 1.89 x 10^-3^ across the four traits (Supplemental Figure S2). The combined significant and top 100 non-significant markers accounted for ∼23% (ranging from 4% to 40%) of the total phenotypic variation (Table S9). The relatively low phenotypic variance explained by these markers reflects the known issue of “missing heritability” in GWAS (Manolio et al., 2009), which is a common phenomenon in which the sum of the effects of significant markers does not fully account for the observed heritability for a trait. However, it has been shown that including markers below the statistical threshold of GWAS minimizes this problem (Yang et al., 2010) and, by incorporating the top 100 non-significant and non-redundant markers, a larger portion of the variability that can be explained by genomic markers was captured. Taken together, these results demonstrate a complex genetic architecture for ear height, plant height, grain moisture, and grain yield in hybrids, and suggests that including additive and dominance effects of markers beyond traditional significance thresholds in simulations may better represent the biologically relevant genetic architecture for these traits.

### 3.2 Traits simulated with the lowest p-value non-redundant GWAS markers have high correlations with empirical data

When simulating traits, multiple parameters must be defined to replicate the genetic architecture of the trait of interest, including the total number of causative variants, the effect size of each causative variant, and the observed genotype for each locus in each individual of the population. To evaluate the number of causative variants required to simulate traits with high correlation to empirical data, many simulation iterations were performed with varying numbers of causative variants. Traits were simulated across multiple environments using the 5, 10, 20, etc. GWAS markers with the lowest mean p-value across environments as the causative variants (Figure 2; Table S10). The effect size for each of these markers was simulated to match the estimated effect size from the GWAS. Concordance of the simulated data with empirical data was measured through Spearman’s Rank Correlation Coefficient to assess rank order, Kolmogorov-Smirnov’s D-statistic to assess distribution shape, and t-test statistic to quantify shifts in distribution means, as well as the log of the F-test statistic to quantify changes in distribution spread. These metrics were measured both within and across environments at the varying number of causative variants.

**Figure 2.**
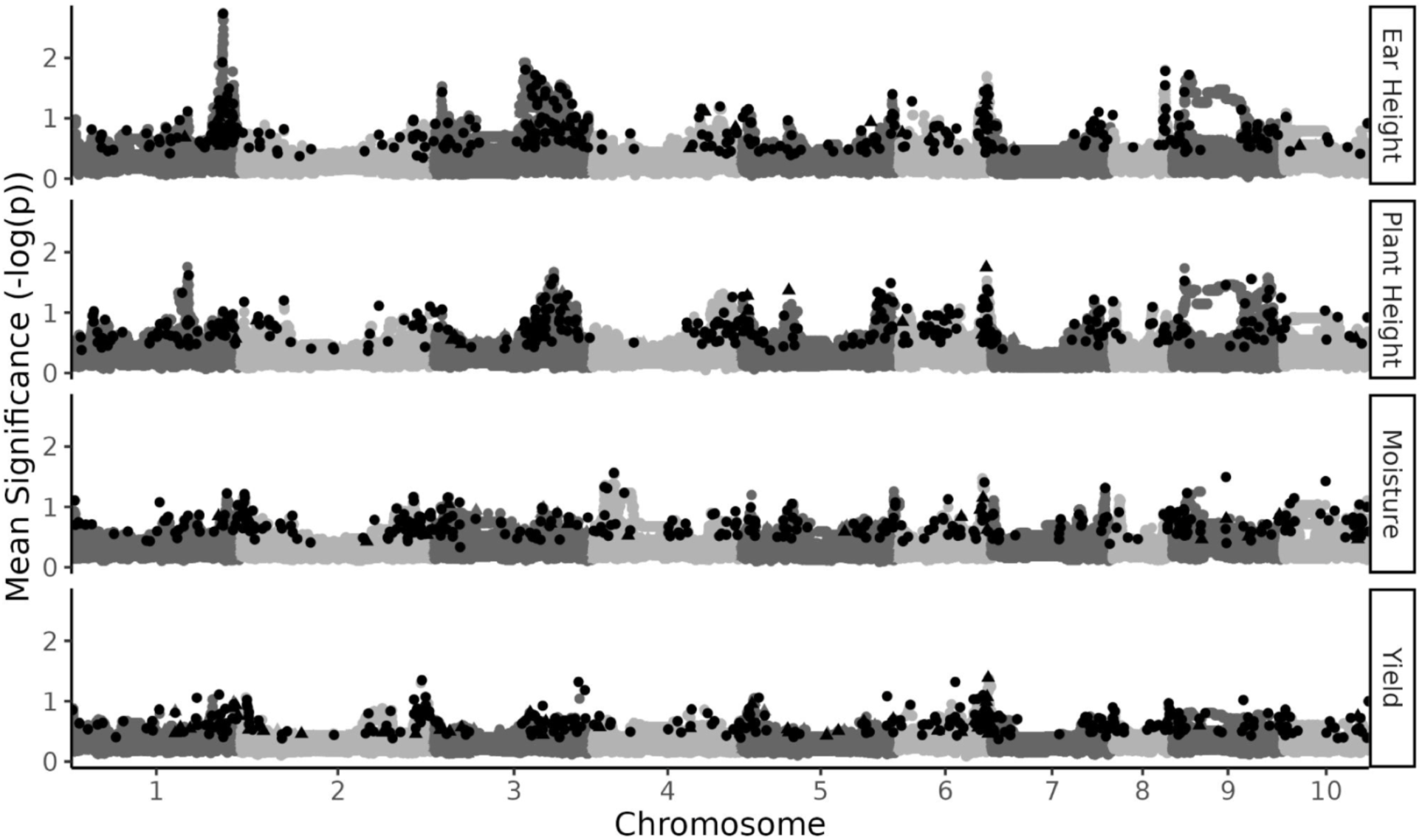
Marker selection from genome wide association study for each trait. Mean marker significance (– log_10_(p)) was calculated across each environment and under both additive and dominance models. The top 350 non-redundant (linkage disequilibrium below background linkage disequilibrium of the population) markers were selected as causative markers for simulation (black) based on the mean association across environments with the trait of interest.

It was hypothesized that concordance between simulated and empirical populations would improve as the number of causative variants increased resulting in increased Spearman’s Rank Correlation Coefficient and all other metrics trending towards zero. Across all four traits, the mean correlations to empirical data within environments increased and the t-test statistic decreased toward zero as the number of causative variants increased (Figure 3A). In contrast to expectation, the distribution dissimilarity increased and the log of F-ratio decreased beyond zero toward more negative values with increased numbers of causative markers. The D-statistic increase indicated that the distribution of the simulated population was becoming less similar to the observed distribution as more causative variants were added. The change in distribution similarity was likely driven by increased range of phenotypic values as the t-test statistic remained near zero (Figure 3A), indicating that the mean of the distribution did not shift, and the log of F-ratio declined to negative values (Figure 3A), indicating that the spread of the simulated values were greater than the observed values.

**Figure 3.**
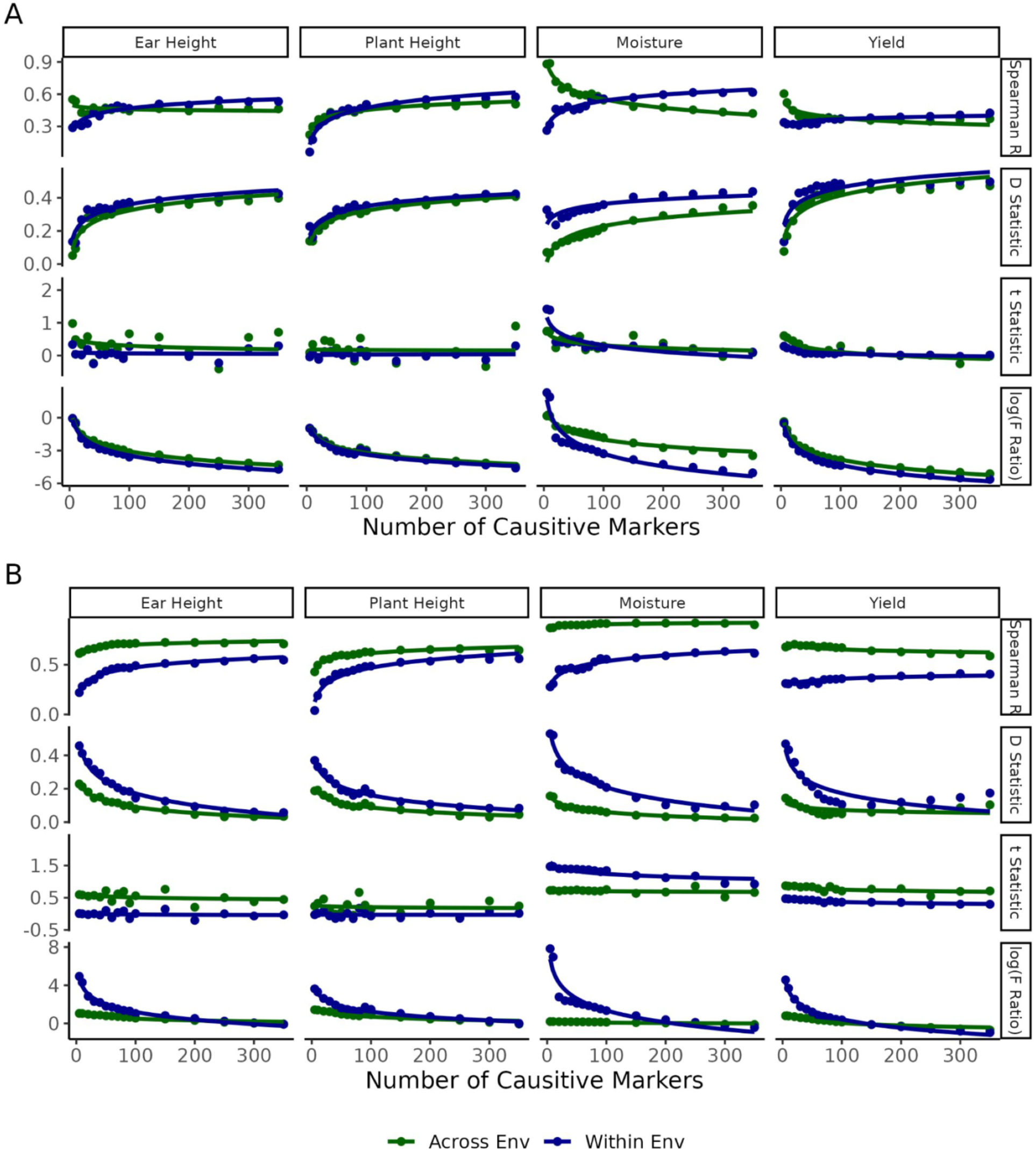
Trait simulation performance over increasing number of causative variants. (A) Average Spearman Rank Correlation Coefficent, Kolmogorov-Smirnov D-statistic, t-test statistic, and log of F-test statistic between simulated and empirical trait values within and across environments from all traits using original GWAS marker effect sizes. (B) Average performance metrics between simulated and empirical data for all traits using one-tenth of the original GWAS marker effect sizes.

The effect of additional causative markers on the concordance between simulated and empirical values was further explored at incremental marker increases, and a plateau in concordance improvement was observed. The mean correlations between simulated and empirical data across traits ranged from 0.249 when the top 5 additive and dominance markers were used to 0.531 when using the top 350 additive and dominant markers. However, the increase was not linear, and a plateau was reached when ∼200 markers were used as causative variants for all traits (mean correlation of 0.505). Similar plateaus at ∼200 markers were reached for the distribution statistics (Figure 3A). These results demonstrate that the genetic architecture of a trait can be sufficiently modeled using ∼200 markers that have the lowest mean p-value across environments from empirical data.

Similar results were observed across environments for each of the concordance metrics in all traits except moisture. The correlation between simulated and empirical moisture values across environments is notably different from the correlation within environments as the number of causative markers increased. This appears to be largely driven by differences in phenotypic values of each environment when few markers are used for simulation (Supplemental Figure S3).

### 3.3 Reducing effect sizes of GWAS hits results in more realistic simulated trait values

The simulated trait data that was generated with the observed effect sizes of the causal variants resulted in unrealistic phenotypic values (Supplemental Figure S4). It is known that GWAS models can overestimate the effect sizes of markers (Goddard et al., 2016), which could contribute to this observation. It was hypothesized that reducing the marker effect size in the simulations could improve the concordance in distribution between simulated and empirical data. To test this hypothesis, the effect sizes from the GWAS were reduced by one order of magnitude.

The mean correlations to empirical data within and across environments still increased as the number of causative variants increased across all four traits. The within-environment correlation values were very similar for traits simulated with reduced and original GWAS effects (Figure 3). In contrast, the correlations across environments were overall higher when simulated with the reduced effect size with a mean increase in correlation of 0.242 (–0.0139 to 0.483). A lesser change in correlation was observed when fewer than 100 causative variants were used to simulate traits with reduced effect sizes (0.0652) compared to original GWAS effects (0.260; Figure 3B).

The impact of reducing the marker effect sizes was more substantial when considering the distribution of the data. The simulated traits with reduced effect size better matched the range of empirical trait values when at least 200 markers were used as causative variants (Figure 3B, Supplemental Figure S5). Using the reduced marker effects, the Kolmogorov-Smirnov D-statistic actually decreased as the number of causative variants increased, indicating increasingly similar distributions (Figure 3B), and aligning with expectations. With this change in marker effect sizes, the t-test statistic remained near zero, indicating minimal difference of population means with the reduced marker effect sizes (Figure 3B). Additionally, the log of the F-test f-ratio decreases towards zero, indicating the variance of the simulated trait distribution was approaching the variance of the empirical data as the number of causative variants increased (Figure 3B). As was seen with the full marker effect size results, a plateau in concordance between simulated and empirical data was reached around 200 causative markers for each trait under all statistics, with the reduced marker effect sizes.

These findings overall demonstrate that reducing the estimated marker effects of genome-wide association studies (GWAS) increases the concordance of simulated data with empirical data. Reducing the effect sizes not only increased correlations across environments, it also increased the similarity of the distribution of the simulated data to the empirical data.

### 3.4 Trait data simulated with reduced marker effect sizes also improves variance partitioning

Another important aspect of trait simulation is to ensure the proper allocation of phenotypic variance into different experimental factors (e.g. genotype, environment, replication within environment, etc.). To assess the accuracy of variance partitioning, ANOVA was performed for each of the simulations and the empirical data and the percent variance explained (PVE) by the experimental factors was calculated (Figure 4). The partitioning of variance for the simulated data generated using the observed marker effect sizes did not have high concordance with the empirical data across the different numbers of markers used in the simulations. At approximately 10 markers used in the simulation, the pattern of trait variance partitioning most closely corresponded to the pattern of trait variance partitioning of their empirical traits (Figure 4). However, using only 10 markers at the original effect sizes resulted in weak correlations between simulated and real phenotypic values (Figure 3A).

**Figure 4.**
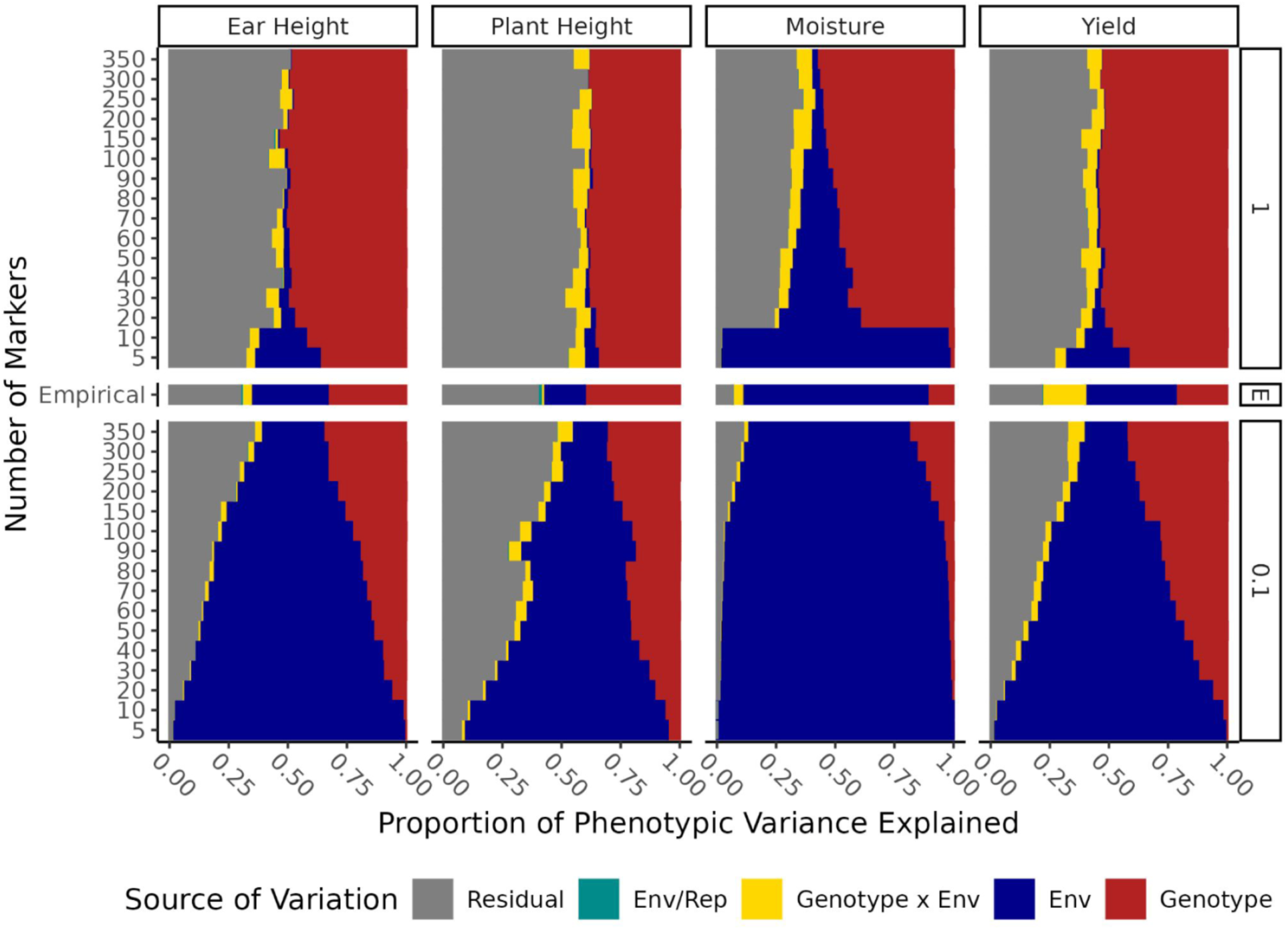
Proportion of phenotypic variance explained in empirical and simulated data. The proportion of phenotypic variance explained was partitioned by genotype, environment, genotype-by-environment, replication-within-environment and residual effects through ANOVA for simulated ear height (cm), plant height (cm), grain moisture (%) and grain yield (bu/ac) with varying number of causative variants (5 through 350) and effect sizes relative to the observed effect sizes from GWAS (0.1 and 1). The proportion of variance explained was compared to the proportion of phenotypic variance explained in the empirical data (E) for all traits.

Variance was also partitioned for simulations based on reduced marker effect sizes given that these simulations improved concordance between simulated and empirical phenotypes. Overall, the concordance with the empirical variance partitioning improved with the reduced marker effect sizes (Figure 4). The number of causative variants with reduced effects that produced the best match between simulated and empirical PVE patterns varied slightly depending on the trait, but using approximately 200 markers provided high concordance in variance partitioning between simulated and empirical trait data (Figure 4). Using fewer markers to simulate traits increased the effects of environment relative to genotype beyond what was observed in the empirical trait data, and using more markers overinflated the effect of genotype relative to environment. These findings suggest a balance is needed in the number of markers and marker effect size used when simulating trait data.

### 3.5 Genomic prediction on simulated traits is useful to test models, but predicted values poorly correlate with empirical traits

In digital breeding, many different genomic prediction models are evaluated with simulated data so that breeders can make informed decisions on the best models to use in their breeding program (Jeon et al., 2023). Thus, to test how well the simulated data generated in this study mirrors the empirical data in evaluating prediction accuracy, genomic prediction was performed for each of the simulated and empirical trait datasets across environments using the same set of 500 randomly selected markers (Table S11). When selecting predictors for the genomic prediction, the causative variants from the GWAS (i.e., the 5 to 350 markers with the lowest p-value) were not masked from selection as predictors, but only 0.4% of predictors, on average, were also causative variants. This approach represents common methods in genomic prediction, where causative markers are not necessarily known, but for highly quantitative traits, are likely to be included by chance.

To obtain a baseline prediction performance to compare to simulation results, genomic estimated breeding values (GEBVs) were determined for empirical data for ear height, plant height, moisture, and yield. Spearman’s Rank Correlation Coefficient, Kolmogorov-Smirnov D-Statistic, t-test statistic, and log of F-test f-ratio were calculated for each trait under an additive and dominance-based model in both a CV1 and CV2 cross-validation scheme. The mean correlation of the additive model was 0.460 (0.171 to 0.636) in CV1 and 0.684 (0.329 to 0.869) in CV2 (Figure 5A). Under the dominance model, the mean correlation was 0.344 (0.139 to 0.474) in CV1 and 0.664 (0.297 to 0.851) in CV2 (Figure 5B).

**Figure 5.**
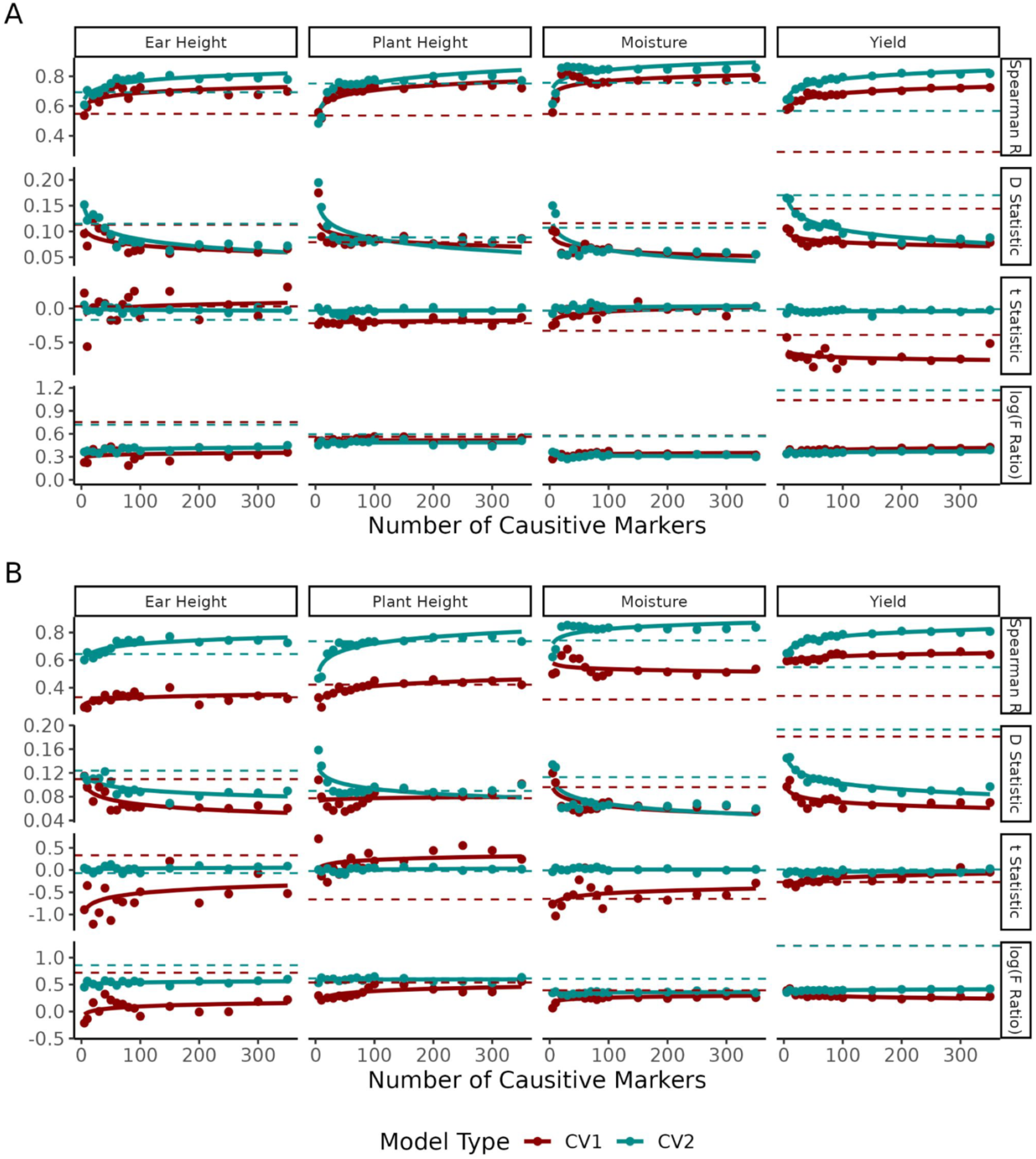
Similarity of genomic prediction values from simulated traits to simulated trait values. (A) Average Spearman Rank Correlation Coefficient, Kolmogorov-Smirnov D-statistic, t-test statistic, and log of F-test statistic between genomic predictions of simulated traits and simulated trait values using an additive model. (B) Average performance metrics between genomic predictions of simulated traits and simulated trait values for all traits using a dominance model. Performance of empirical models trained under similar parameters are shown with dashed lines.

To assess the parallels of the simulated trait data for utility in making breeding decisions, GEBVs of simulated data were compared to the phenotypes simulated with reduced marker effect sizes. A high degree of concordance was observed between genomic prediction of simulated data with the simulated phenotypes, which continued to improve as the number of causative markers increased in the simulation under both additive (Figure 5A) and dominance models (Figure 5B). The mean correlation of simulated trait GEBVs to the simulated traits was 0.613 (0.200 to 0.834) at 200 causative markers in CV1 and 0.803 (0.660 to 0.909) at 200 causative markers in CV2. It is notable that the concordance between GEBVs of simulated data and simulated trait values was better than that observed between GEBVs of empirical data and observed empirical data for all traits in all metrics except the t-test statistic. This suggests that while the mean values of GEBVs shift slightly when trained on simulated data, the rank and distribution shape tend to be representative. This high concordance of rank and distribution shape could be due in part to the simulated traits more closely aligning with the assumptions of the infinitesimal model that the genomic prediction model is based on.

The GEBVs of the simulated data were next compared to the true empirical data to assess concordance across multiple levels of inference (i.e., simulation of trait data and then prediction from simulated data) and utility as a digital twin. When trait data was simulated using 200 causative markers, the mean correlation observed across traits ranged from 0.344 to 0.529, the mean D-statistic ranged from 0.120 to 0.156, the mean t-test statistic ranged from –0.325 to 1.66, and the mean log of f-ratio ranged from –0.0973 to 1.32. The concordance metrics were very similar across both additive and dominance models and CV1 and CV2 cross-validation schemes. The primary difference was a decrease in correlation performance in the CV1 assessment for ear height, plant height, and moisture when using a dominance model. Increasing the number of causative variants led to a small increase in the mean correlation within environments for traits like yield, moderate increases for ear height and moisture, and significant increases for plant height in both additive (Figure 6A) and dominance models (Figure 6B). Although this pattern aligns with that between the simulated and empirical values (Figure 3), the absolute correlation values between GEBVs of simulation values and empirical data are lower than the correlations between simulated and empirical data (e.g., ∼0.451 and ∼0.505, respectively, across all traits with top 200 causative variants and reduced effect sizes; Figure 3, Figure 6). Additionally, as the number of causative markers was increased, the distribution of simulated trait GEBVs became more similar to the distribution of empirical data with the Kolmogorov-Smirnov D-statistic declined, t-test statistic remained near zero, and the log of the F-test f-ratio approached zero in both CV1 and CV2 schemes (Figure 6A, Figure 6B). Moisture was the exception to this trend. For moisture, a considerable shift in mean was observed regardless of the number of causative variants (Figure 6A, Figure 6B). Overall, these results show that as the number of causative markers increased in the simulations, the genomic predictions of simulated traits became more concordant with observed phenotypes.

**Figure 6.**
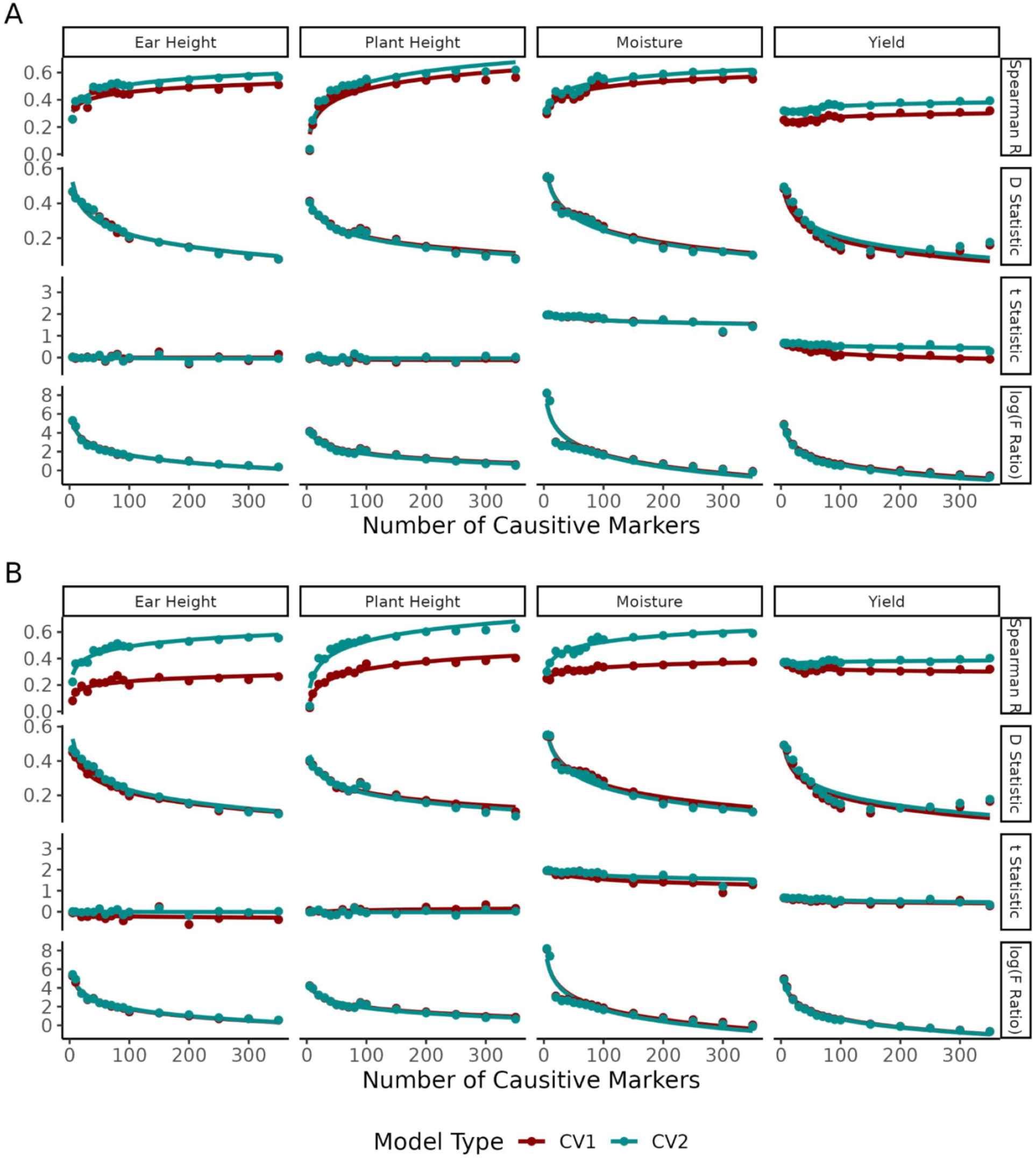
Similarity of genomic prediction values from simulated traits to empirical trait values. (A) Average Spearman Rank Correlation Coefficent, Kolmogorov-Smirnov D-statistic, t-test statistic, and log of F-test statistic between genomic predictions of simulated traits and empirical trait values using an additive model. (B) Average performance metrics between genomic predictions of simulated traits and empirical data for all traits using a dominance model.

In addition to comparing GEBVs of simulated traits to empirical traits, the correlation between GEBVs of simulated traits and GEBVs of empirical traits was assessed to determine the utility of simulating early breeding stages where selections may be made solely on genetic information. GEBVs of simulated traits correlated to GEBVs of empirical traits with a mean Spearman’s Rank Correlation Coefficient of 0.688 (range: 0.503 – 0.801) when 200 markers were used in CV2 validation (Figure 7). As the number of causative variants increased under both additive (Figure 7A) and dominance models (Figure 7B), a higher concordance, both in terms of correlations and converging distributions, was observed. The only notable exception to this was yield, in which a divergence of distributions was observed when the causative markers increased beyond 100 markers.

**Figure 7.**
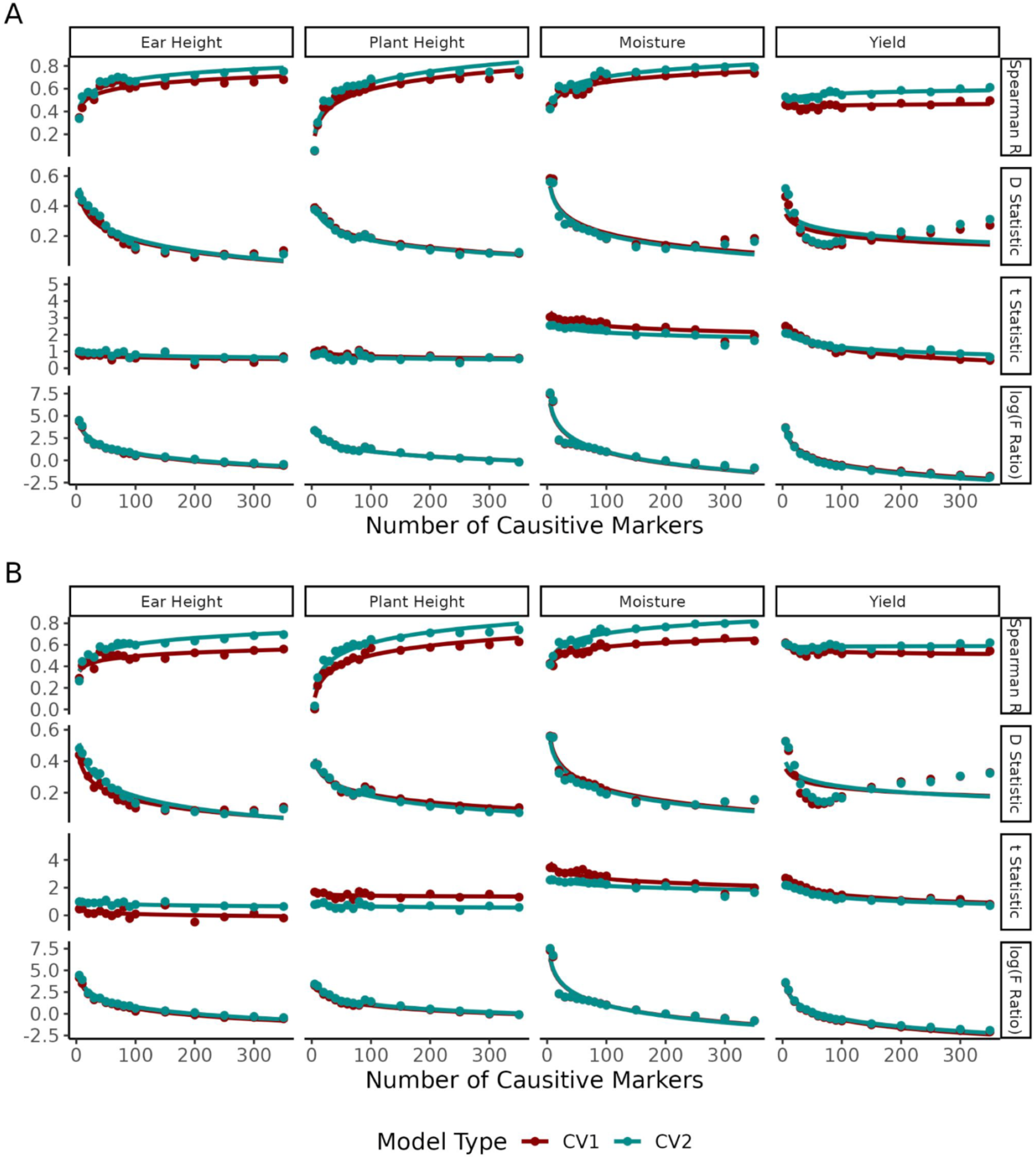
Similarity between genomic predicted values from simulations and genomic predicted values of empirical traits. (A) Average Spearman Rank Correlation Coefficent, Kolmogorov-Smirnov D-statistic, t-test statistic, and log of F-test statistic between genomic predictions of simulated traits and genomic predictions of empirical trait values using an additive model. (B) Average performance metrics between genomic predictions of simulated traits and genomic predictions of empirical data for all traits using a dominance model.

## 4 CONCLUSION

This study evaluated different methods for simulating traits across multiple environments for common real-world agronomic traits, including those that are highly influenced by environmental and genotype-by-environment effects. The results of this study showed that accounting for the genetic architecture of traits, particularly beyond traditional significance thresholds, in trait simulations will result in simulated data that is in high concordance with observed empirical traits. Utilizing at least 200 markers with the highest association yielded the best concordance with empirical data means and distributions in this study. Reducing the estimated marker effects also produced more realistic distributions of phenotypic values and more realistic partitioning of phenotypic variances. Finally, the predicted phenotypic values (GEBVs) from simulated data were moderately correlated with the respective empirical trait data, indicating utility for testing different breeding strategies. Simulating agronomic traits that match real-world data is a valuable tool for plant breeders to create a digital version of their breeding program that is relevant to their specific germplasm and not based on random genetic architectures. These digital breeding programs can be used to improve resource allocation, evaluate and enhance germplasm throughout a breeding program by assessing many potential genetic recombinants *in silico*, augment existing phenotypic data to provide the sample sizes needed for training machine learning models for genomic prediction, and to assess the impact of forecasted climate shifts on germplasm. The ability to accurately assess germplasm *in silico* has the potential to greatly reduce the time and cost associated with plant breeding.

## SUPPLEMENTAL MATERIAL

Supplemental Figure 1. Significant markers from GWAS across traits, environments, and models.

Supplemental Figure 2. Distribution of p-values for each GWAS performed.

Supplemental Figure 3. Correlation between simulated trait values and empirical trait values with full marker effects.

Supplemental Figure 4. Distribution of empirical and simulated trait values with an increasing number of causative variants.

Supplemental Figure 5. Correlation between simulated trait values and empirical trait values with reduced marker effects.

Supplemental Table 1. Hybrids developed from a diallele cross between 333 recombinant inbred lines.

Supplemental Table 2. Growing environment information for empirical data collection.

Supplemental Table 3. Phenotypic measurements of 400 hybrids across 11 environments

Supplemental Table 4. Weather variables collected from the EnvRtype package and their definitions

Supplemental Table 5. Weather data collected for each site during the 2019 and 2020 growing seasons.

Supplemental Table 6. Variance-covariance relatedness matrix of environments based on 17 weather parameters in ten-day intervals throughout the growing season.

Supplemental Table 7. Proportion of variance explained and significance of each source of variation.

Supplemental Table 8. Significant, non-redundant markers for each trait in each environment from each gwas model

Supplemental Table 9. Proportion of phenotypic variance explained by genetic markers

Supplemental Table 10. Markers chosen for simulation. Markers were chosen as the top 350 non-redundant markers with the highest mean association with the trait of interest.

Supplemental Table 11. Markers chosen for genomic prediction.

## DATA AND CODE AVAILABILITY

The data that supports the findings of this study are available in the supplementary material of this article and the data repository https://hdl.handle.net/11299/272874. Raw phenotypic trait data is available in Table S3, environmental data is available in Table S5, and marker data were downloaded from https://doi.org/10.13020/atq4-1b58 (Della Coletta et al., 2023). All code used in this study is publicly available on GitHub at https://github.com/HirschLabUMN/empirical_simulation.

## AUTHOR CONTRIBUTIONS

CNH, AEL, MOB, SBF conceived this experiment and collected the empirical data. MJB, RDC conducted the data analysis. MJB, RDC, CNH wrote the original draft. All co-authors edited and approved the final manuscript.

## ACKNOWLEDGEMENTS

We thank Bayer Crop Science, Corteva Agriscience, Syngenta, and Beck’s Hybrid Corn Seed for providing in-kind support through growing locations for the multi-environment yield trial data used in this study and DOW AgroScience (now Corteva Agriscience) for providing in-kind support through the custom Illumina Infinium 20k SNP chip. We thank the Minnesota Supercomputing Institute at the University of Minnesota (http://www.msi.umn.edu) for providing resources that contributed to the research results reported in this article. This work was supported by the United States Department of Agriculture (2018-67013-27571 and 2024-67013-42588) and the Minnesota Agricultural Experiment Station. RDC was supported by the University of Minnesota MnDRIVE Global Food Ventures Graduate Fellowship and the University of Minnesota Doctoral Dissertation Fellowship. MJB was supported by the University of Minnesota MnDRIVE Informatics Institute Fellowship and the University of Minnesota Doctoral Dissertation Fellowship.

## ABBREVIATIONS

BLUE: best linear unbiased estimate
CV1: cross validation 1
CV2: cross validation 2
GBLUP: genomic best linear unbiased prediction
GEBV: genomic estimated breeding value
GWAS: genome-wide association study
LD: linkage disequilibrium
MLM: mixed linear model
PCA: principal component analysis
PVE: proportion of variance explained
QTL: quantitative trait loci
SNP: single nucleotide polymorphism
SV: structural variant

