## Supplemental Figures for "Optimizing population simulations to accurately parallel empirical data for digital breeding"

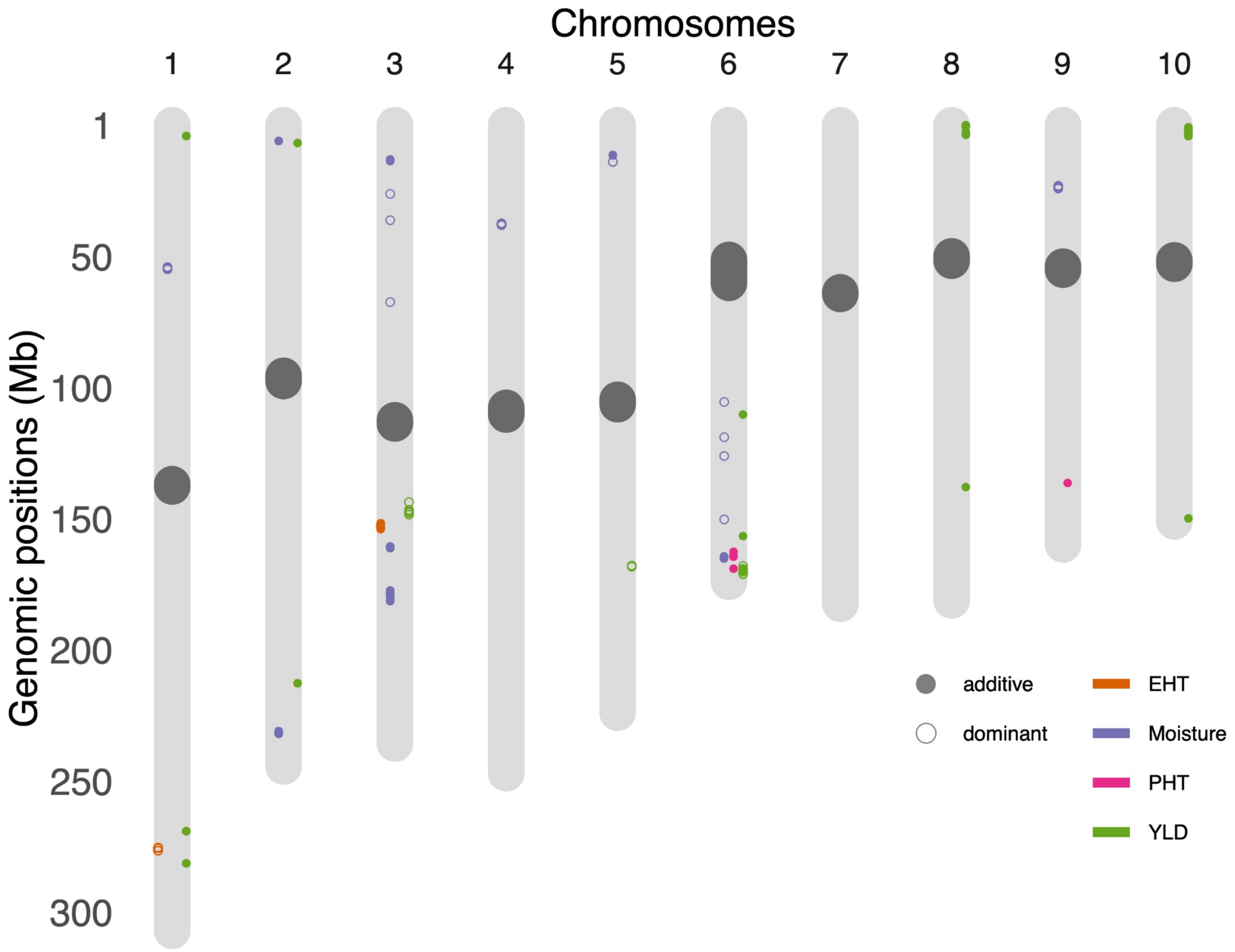


**Supplemental Figure 1**. **Significant markers from GWAS across traits, environments, and models.** Results of genome-wide association studies (GWAS) for ear height (EHT), plant height (PHT), grain moisture (Moisture) and grain yield (YLD) with additive and dominance models. Karyoplot shows the genomic location of markers statistically significantly associated with each trait.

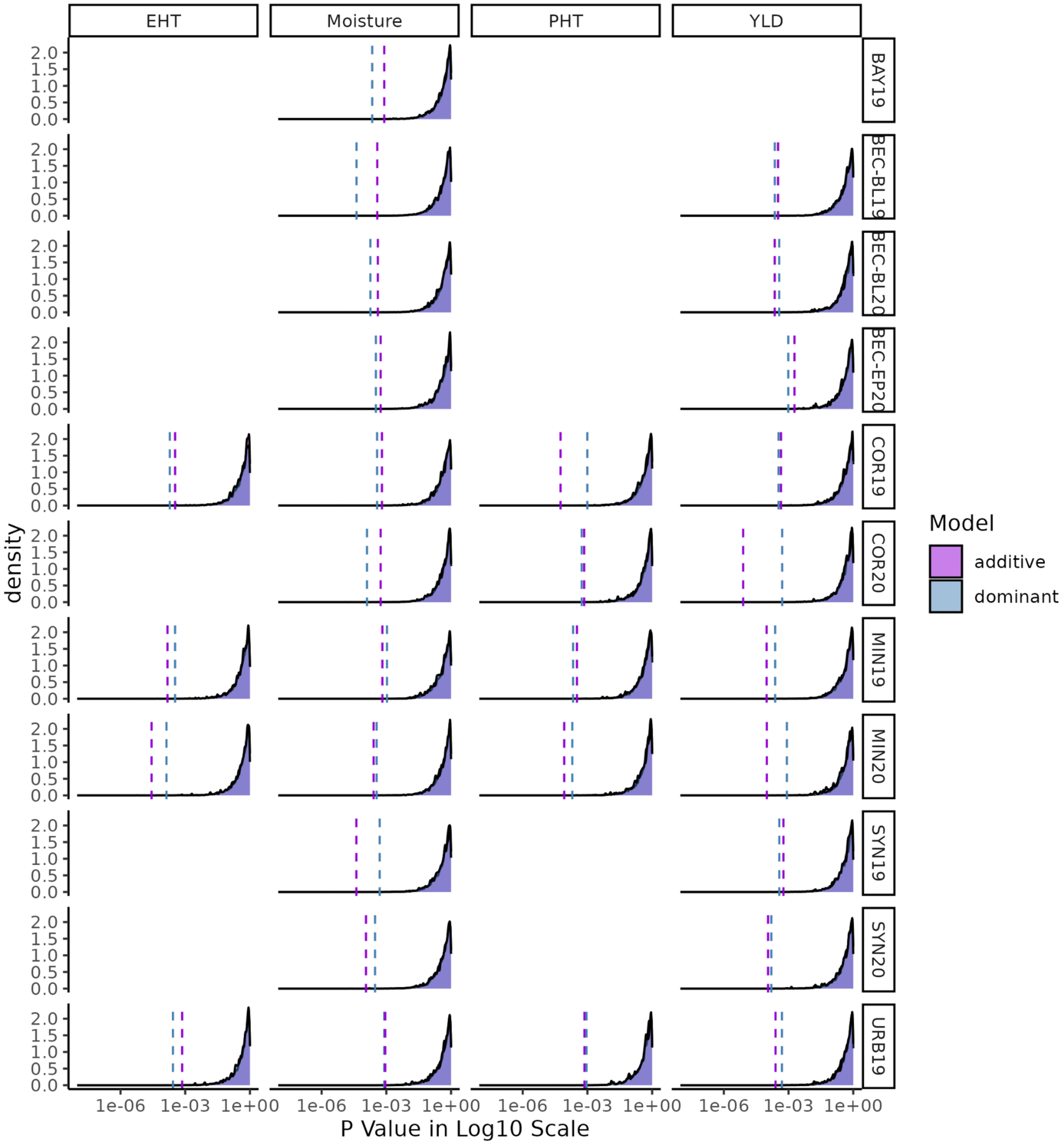


**Supplemental Figure 2**. **Distribution of p-values for each GWAS performed.** P-value results of genome-wide association studies (GWAS) for ear height (EHT), plant height (PHT), grain moisture (Moisture) and grain yield (YLD) with additive and dominance models. Vertical dashed lines indicate the p-value of the hundredth non-significant p-value for each GWAS.

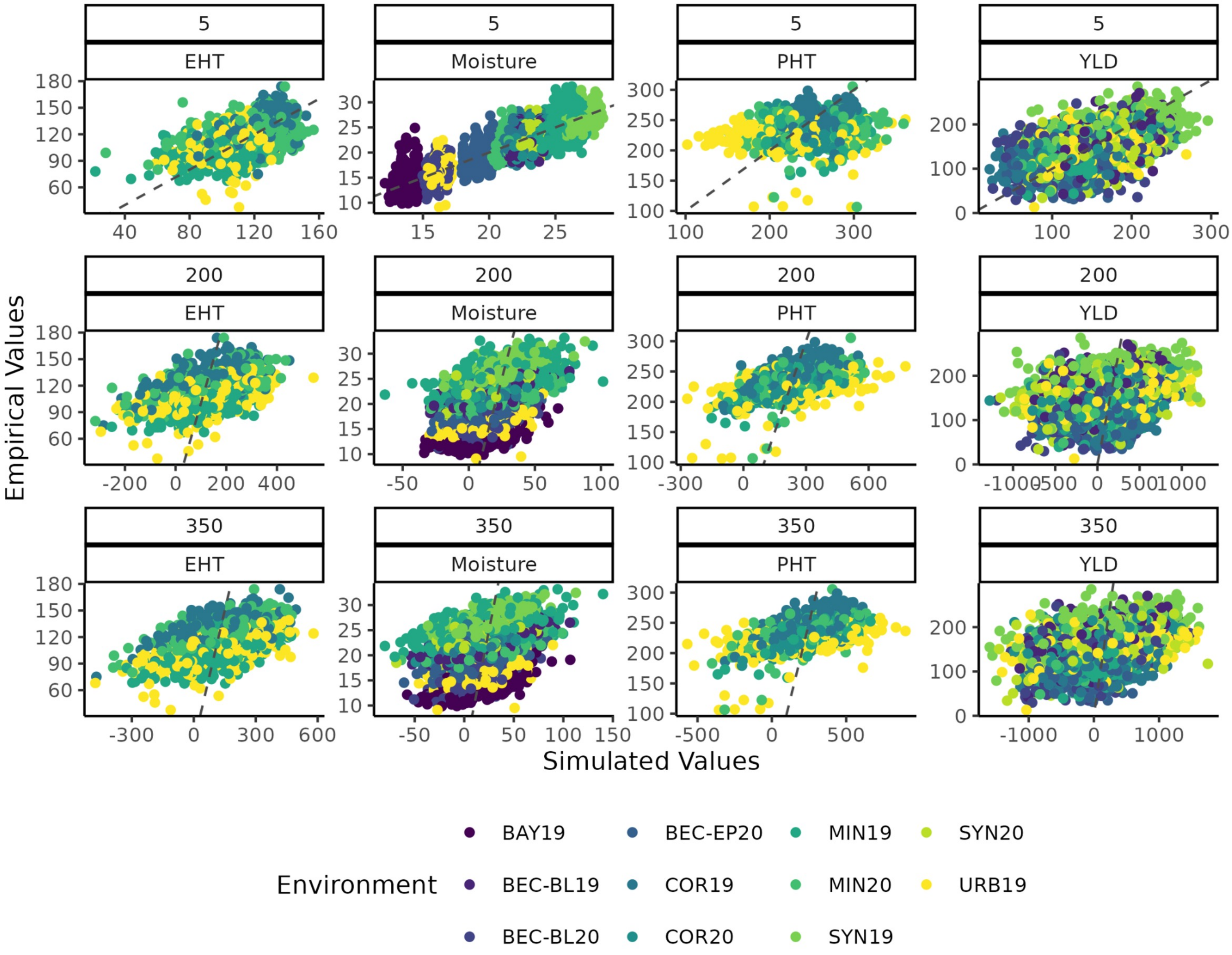


**Supplemental Figure 3**. **Correlation between simulated trait values and empirical trait values with full marker effects.** Colors correspond to environments, and plots based on 5, 200, and 350 causative variants are shown to illustrate the extremes. The gray dashed line represents a perfect correlation with a slope of one and intercept of zero.

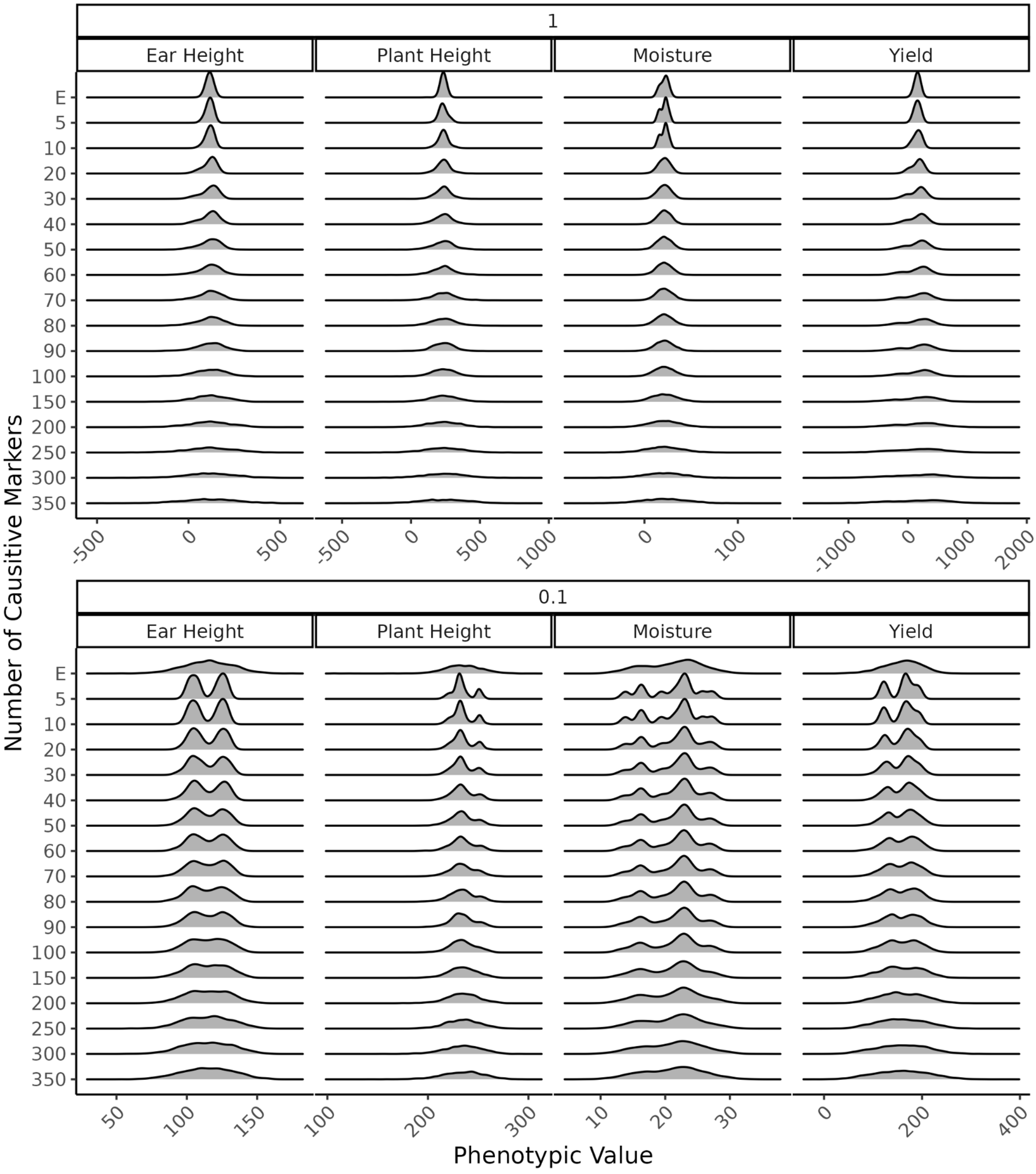


**Supplemental Figure 4**. **Distribution of empirical and simulated trait values with an increasing number of causative variants.** Density plots of empirical (E) and simulated trait values with increasing number of causative variants (5 to 350). Distributions are shown for simulations that used GWAS reported marker effect size (1) and reduced marker effect size (0.1).

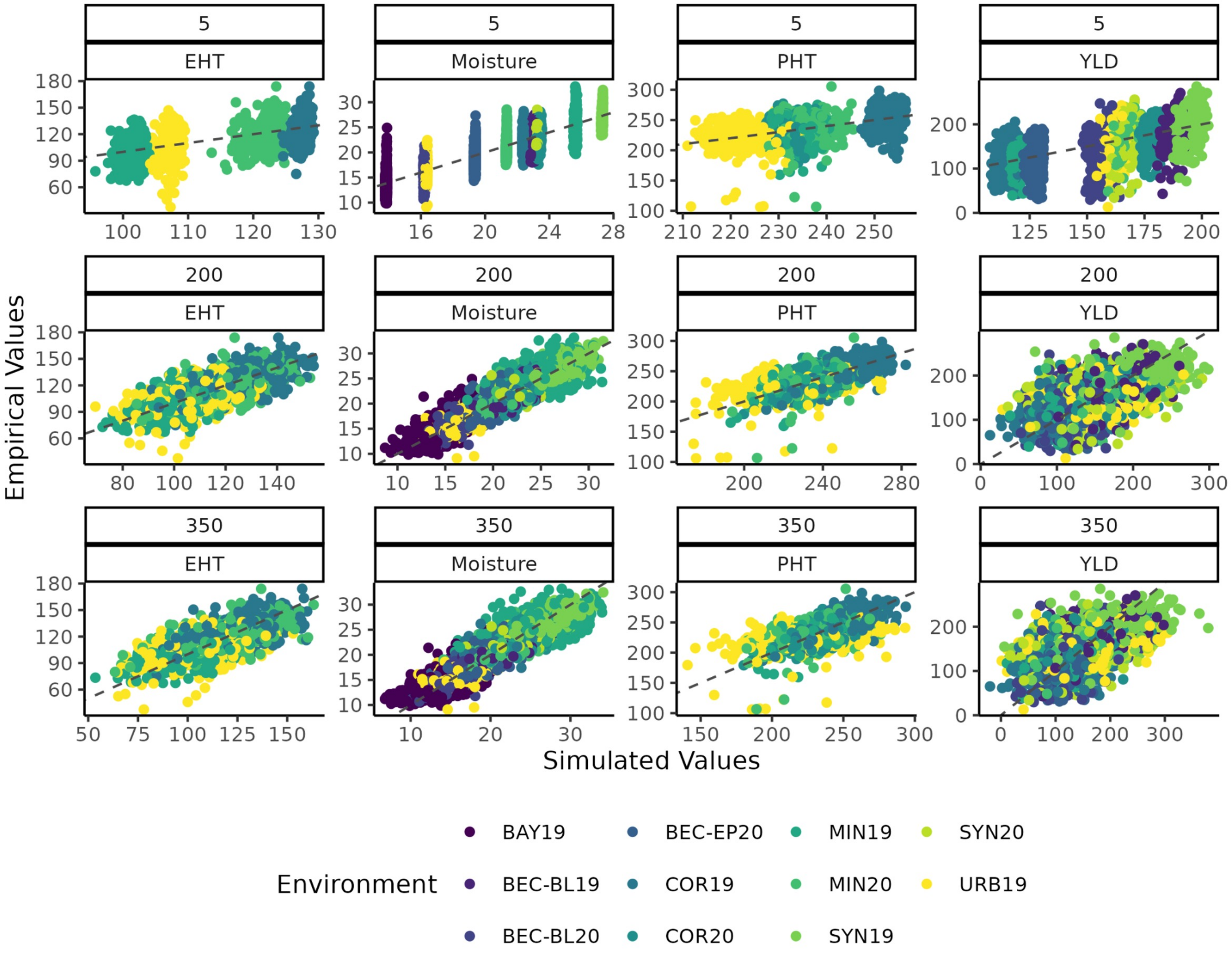


**Supplemental Figure 5**. **Correlation between simulated trait values and empirical trait values with reduced marker effects.** Colors correspond to environments, and plots based on 5, 200, and 350 causative variants are shown to illustrate the extremes. The gray dashed line represents a perfect correlation with a slope of one and intercept of zero.
